# *Drosophila* larval light avoidance is negatively regulated by temperature through two pairs of central brain neurons

**DOI:** 10.1101/2020.11.24.395848

**Authors:** Jie Wang, Weiqiao Zhao, Qianhui Zhao, Jinrun Zhou, Xinhang Li, Yinhui He, Zhefeng Gong

## Abstract

Animal’s innate avoidance behavior is crucial for its survival. It subjects to modulation by environmental conditions in addition to the commanding sensorimotor transformation pathway. Although much has been known about the commanding neural basis, relatively less is known about how innate avoidance behavior is shaped by external conditions. Here in this paper, we report that *Drosophila* larvae showed stronger light avoidance at lower temperatures than at higher temperatures. Such negative regulation of light avoidance by temperature was abolished by blocking two pairs of central brain neurons, ACLP^R60F09^ neurons, that were responsive to both light and temperature change, including cooling and warming. ACLP^R60F09^ neurons could be excited by *pdf*-LaN neurons in the visual pathway. On the downstream side, they could inhibit the CLPN^R82B09^ neurons that command light induced reorientation behavior. Compared with at warm temperature, ACLP^R60F09^ neurons’ response to light was decreased at cool temperature so that the inhibition on CLPN^R82B09^ neurons was relieved and the light induced avoidance was enhanced. Our result proposed a neural mechanism underlying cross-modal modulation of animal innate avoidance behavior.

## Introduction

The neural mechanism underlying animal preference behavior has been intensively studied as it reflects the most common form of sensorimotor transformation (Glimcher, 2003; Poeppl et al., 2016; Song and Lee, 2018; Salamone et al., 2018; Lowenstein and Velazquez-Ulloa, 2018; Marachlian et al., 2018). For animal innate preference behaviors, in addition to the commanding internal neural mechanism, they can also be modified by external environmental conditions. In *Drosophila* for example, preference for food containing different concentration of sugar was biased by hardness of food (Jeong et al., 2016). Adult females choose site of egg laying according to the interplay between sweet taste and mechanical feeling of hardness (Wu et al., 2019). Flies’ humid preference was reported to be affected by temperature sensation while rapid heat avoidance was negatively related to environmental humidity on the other hand (Frank et al., 2017). Also, temperature preference in adult fly was positively influenced by environmental light conditions (Head et al.,2015). Such environmental modulation of innate preference behavior may facilitate the animals’ response to those environmental conditions under which certain external stimuli usually co-exist.

Comparatively, environmental modulation of *Drosophila* light preference has received less investigation. *Drosophila* avoids light and prefers darkness at larval stage (Keene et al., 2011; Keene and Sprecher 2012). In *Drosophila* larva, the anteriorly located larval photoreceptors, Bolwig’s organs, which sense low- or intermediate-intensity light and are required for light avoidance response (Mazzoni et al., 2005; Keene et al. 2011; Keene and Sprecher, 2012; Humberg and Sprecher, 2017; Humberg et al., 2018). Bolwig’s organs project their axons into the larval optic neuropil (LON) and form synapses with downstream neurons such as *pdf*-expressing lateral neurons (*pdf-*LaN) Larderet et al., 2017), the fifth lateral neuron (5^th^-LN) (Keene et al. 2011) and the PVL09 neurons (Humberg et al., 2018), with the latter two reported to be required for larval light avoidance. Downstream to these neurons includes a neural pathway consisting of PTTH neurons (Gong et al., 2010), EH neurons and Tdc2 motor neurons that commands light induced deceleration response (Gong et al., 2019), and another pathway including LRIN^R13B07^ neurons and CLPN^R82B09^ neurons that mediates light induced reorientation (Zhao, et al., 2019).

As light is closely related to temperature such as in daily or seasonal cycle, it is assumable that *Drosophila* larval light avoidance is under the impact of temperature. Indeed, we could observe stronger light avoidance at lower temperatures than at higher temperatures. In *Drosophila* larva cold-sensing neurons are anteriorly located with cell bodies in the dorsal organ ganglion (DOG) and the terminal organ ganglion (TOG) (Liu et al. 2003; Klein et al., 2015; Li and Gong, 2015; Ni et al., 2016; Li and Gong, 2017). They send sensory dendrites to terminal organs and dorsal organs in the very anterior tip and axonal projections to central brain. DOG neurons sense cold stimuli through mediation of ionotropic receptors IR21a and IR25a (Ni et al., 2016). On the other hand, neurons expressing TRPA1 and *painless* were found to sense innocuous warmth and nociceptive heat in *Drosophila* larva (Barbagallo and Garrity, 2015; Rosenzweig et al., 2005; Tracey et al., 2003). Interestingly, the light receptor protein Rh5 and Rh6 were also reported to regulate larval temperature preference behavior (Sokabe et al., 2016). There is relatively little study on how cold and warmth signal is processed in downstream neurons (Li and Gong, 2015).

In this study, we report that *Drosophila* larval light avoidance was negatively regulated by temperature. Such thermal regulation of light avoidance was abolished by blocking two pairs of ACLP^R60F09^ neurons that could sense both light stimulation and temperature changes including cooling and warming. At lower temperature, the light induced response in ACLP^R60F09^ neurons was repressed so that their inhibition on downstream CLPN^R82B09^ neurons that are known to mediate light induced reorientation response was relieved. Our results reveal a neural mechanism underlying modulation of light evoked animal avoidance behavior by environmental temperature.

## Results

### 1. Drosophila larval light avoidance was negatively regulated by temperature

We investigated the effect of temperature on larval light avoidance by testing 3^rd^ instar larval light preference using a simple light/dark choice assay (Gong et al., 2010; Zhao et al., 2019) with white light intensity of 550 lux (about 23.3 μW/mm^2^) for 10 minutes at temperatures ranging from 15 °C to 27 °C using a wildtype strain of WT-CS. In this range, larval light avoidance was promoted as temperature decreased (Fig. 1a). Furthermore, we asked if the behavioral output was stable over longer time of testing. We tested 3^rd^ instar larval light avoidance of another commonly used control strain *w^1118^* at 18 °C and 27 °C for 5-, 10-, 15- and 20 minutes respectively. In all cases, larvae always showed a higher preference for darkness at a cool temperature of 18 °C than at a warm temperature of 27 °C (Fig. 1b). These results together showed that larval light avoidance was negatively regulated by temperature.

**Fig. 1.**
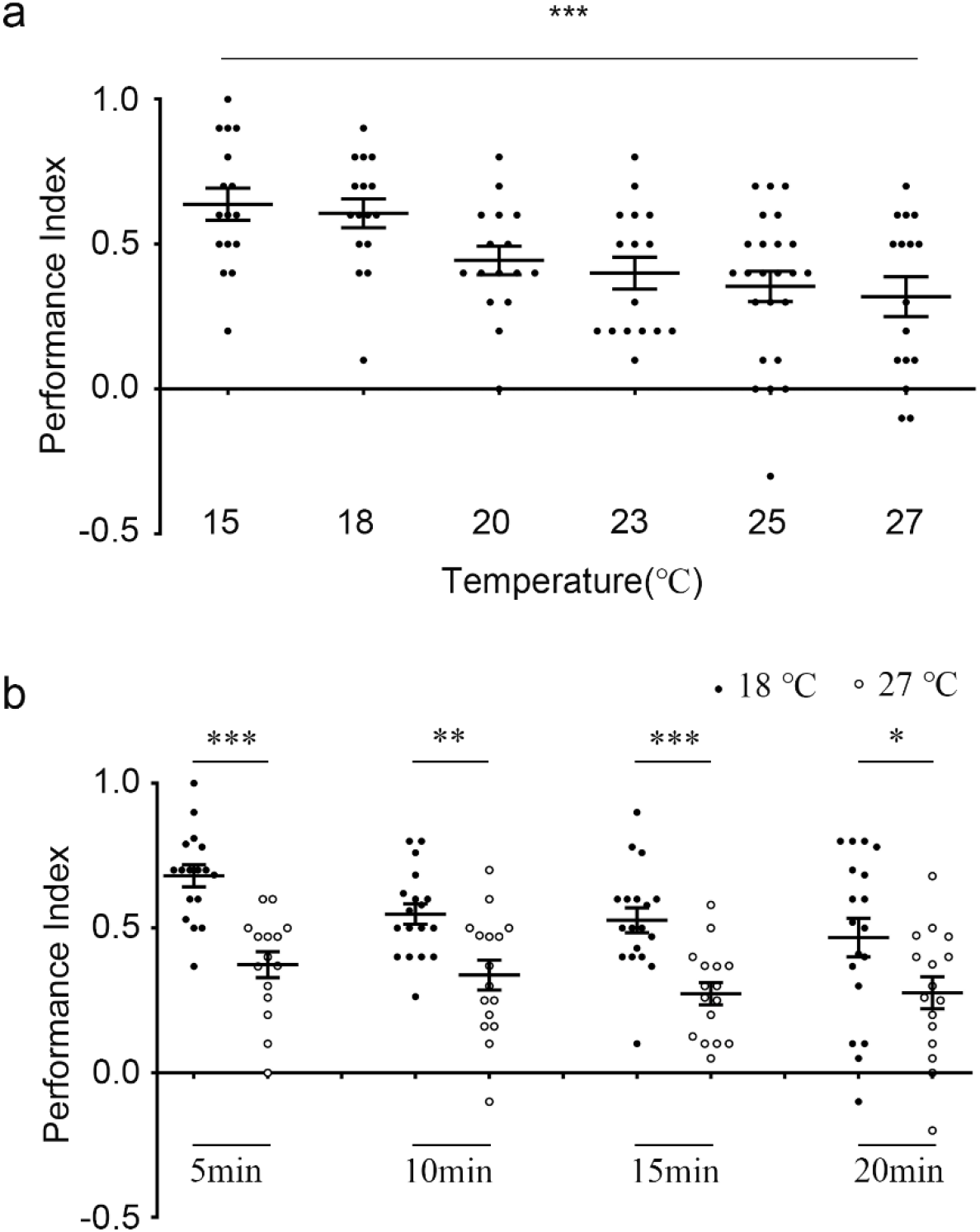
Negative regulation of light avoidance by temperature in *Drosophila* Larvae. (a) WT-CS larvae show stronger light avoidance at lower temperature in a light/dark choice assay for 10 minutes. *** *P* < 0.001, one-way ANOVA followed by post hoc Tukey’s multiple comparison test; n = 16 for all temperatures except that n = 24 for 25 °C. (b) *w^1118^* larvae always show stronger light avoidance at 18 °C than at 27 °C at different time points of test. *** *P* < 0.001, ** *P* < 0.01, * *P* < 0.05, unpaired *t*-test; n = 17, 15, 18, 16, 18, 16, 18, and 16 from left to right). White light of 550 lux (23.3 μW/mm^2^) was used in both (a) and (b). Error bars, SEMs.

### 2. R60F09-GAL4 labeled neurons were putatively required for regulation of light avoidance by temperature

In order to explore the neural basis of lower temperature enhancement of larval light avoidance, we screened for candidate neurons by using Gal4 strains to drive the expression of tetanus toxin (TNTG) to block the activity of neurons and comparing larval light avoidance performance to 170 lux white light at 18 °C and 27 °C. The difference between 18 °C and 27 °C seen in control lines disappeared when TNTG was expressed by *R60F09-GAL4* (Fig. 2a). In larval central nervous system, *R60F09-GAL4* marks three pairs of neurons in the brain hemispheres and three pairs of neurons in the suboesophageal zone (SEZ) (Fig. 2b). We used flp-out technology (Zhao et al., 2019) to reveal single neuronal morphology in the brain and found that the three pairs of neurons in the brain were divided into two categories: in the first category, each of the two pairs of neurons has dendrites and cell body ipsilaterally located in anterior brain hemisphere and an axon projecting contralaterally to the other hemisphere (Fig. 2c, e); in the second category, each of the third pair of neurons has cell body and dendritic arborizations as well as axonal arborizations all in the same brain hemisphere (Fig. 2d). For convenience, we named neurons in the first category ACLP^R60F09^ neurons (anterior contralateral projecting neurons). Thus, *R60F09-GAL4* labeled neurons were potentially required for the regulation of larval light avoidance by temperature.

**Fig. 2.**
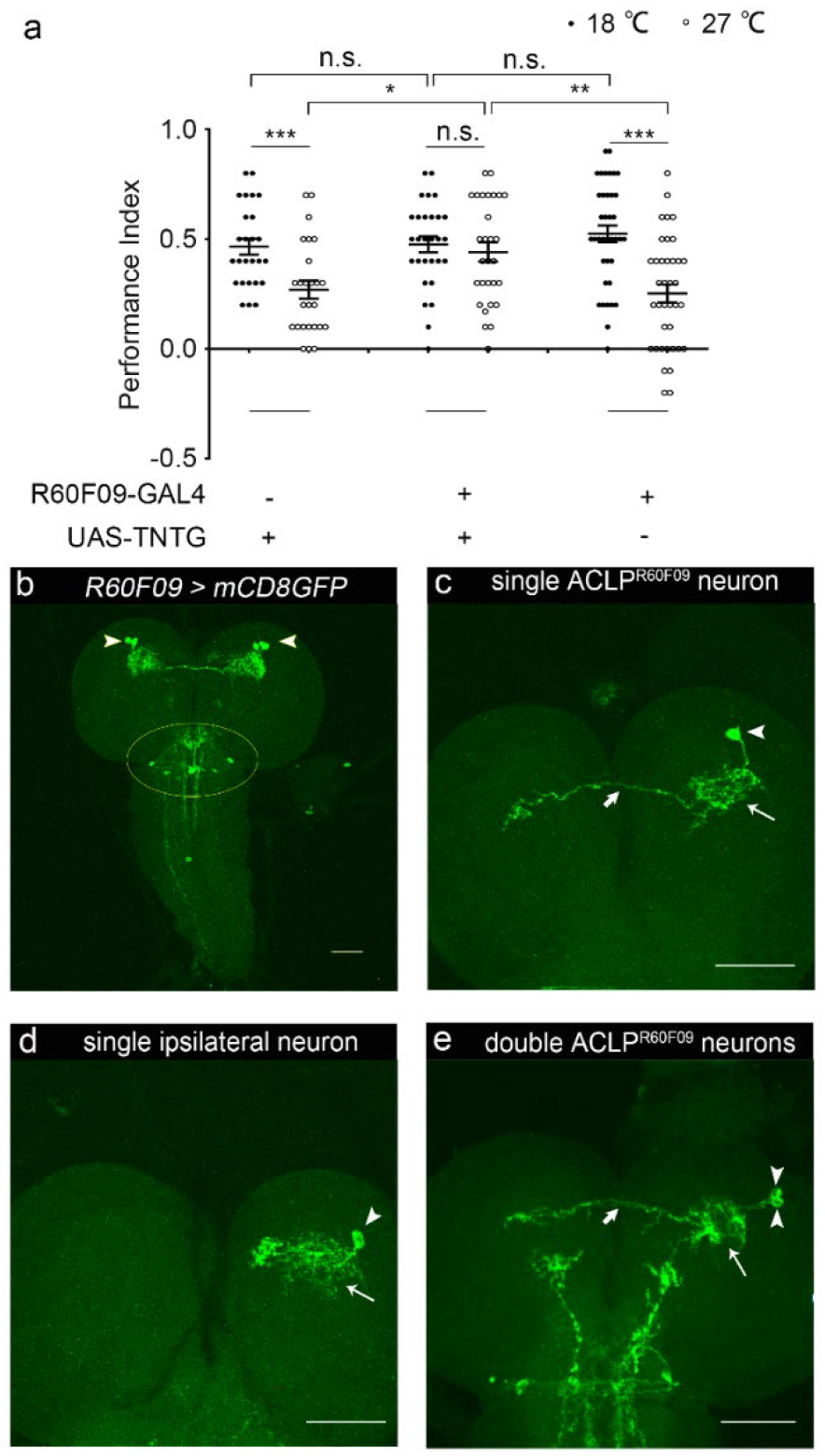
ACLP^R60F09^ neurons participate in thermal regulation of larval light avoidance. (a) Blocking ACLP^R60F09^ neurons using TNTG abolished difference in light avoidance at cool and warm temperatures in a light/dark preference assay using white light at intensity of 170 lux. n.s. *P* > 0.05, *** *P* < 0.001, unpaired t-test; n = 26, 26, 29, 29, 38 and 38 from left to right. Note that performance index of larvae with ACLP^R60F09^ neurons blocked using TNTG is higher than in control *Drosophila* larvae at 27 °C (* *P* < 0.05, \*\**P* < 0.01, one-way ANOVA followed by post hoc Tukey’s multiple comparison test; n = 26, 29 and 38 from left to right). Error bars, SEMs. (b) Expression of *R60F09-GAL4* in larval central nervous system. Arrowheads indicate the ACLP^R60F09^ neurons. Yellow circle shows the neurons in SEZ. (c, d) Morphology of single *R60F09-GAL4* neurons in brain hemispheres, including ACLP^R60F09^ neuron (c) and the 3^rd^ neuron lacking contralateral projection (d). (e) Morphology of two ACLP^R60F09^ neurons located in the same brain hemisphere. Arrowheads point to the cell bodies. Long thin arrows point to dendrites. Short thick arrows point to axonal projections. Scale bars, 50 μm.

### 3. ACLP^R60F09^ neurons were responsive to both light and temperature changes

We proposed that thermal sensation might affect larval light avoidance by modulating the neurons in the underlying pathway. If so, the neurons crucial for the cross-modality sensory integration should be responsive to both light and temperature change. We then tested this hypothesis using calcium imaging. Upon light stimulation, the two ACLP^R60F09^ neurons in the same brain hemisphere showed obvious increase in calcium signal as indicated by GCAMP, with average peak response of about 10% increase in fluorescent intensity, while the third neuron in brain hemisphere as well as neurons in SEZ did not show obvious response (Fig. 3a and 3b, Supplemental Figure 1). Treating the neurons with 20 μM tetrodotoxin (TTX) (Pirez et al., 2013; Luo et al., 2017), a voltage-gated Na^+^ channels antagonist, could completely abolish the response, suggesting that the ACLP^R60F09^ neurons do not sense light by themselves but receive light signal from other neurons (Fig. 3a). Therefore, we focused only on the two pairs of ACLP^R60F09^ neurons in following experiments. We next tested the response of ACLP^R60F09^ neurons to temperature change. As shown in Fig 2a, blocking *R60F09-GAL4* labeled neurons appeared to promote larval light avoidance at 27 °C but not at 18 °C, suggesting that it could be the temperature rise that inhibited light avoidance. We first tested ACLP^R60F09^ neurons’ response to temperature rise. As shown in Fig 3c and 3d, temperature rise monotonically from 18 °C to 27 °C within 30 seconds using a temperature controller led to a fast decrease in calcium signal by about 30% in peak response, suggesting that ACLP^R60F09^ neurons were repressed by temperature rise. Application of 20 μM tetrodotoxin (TTX) could largely remove the inhibitory response, suggesting that the thermal response originated from other neurons but not the ACLP^R60F09^ neurons themselves. We then wanted to know if temperature drop could produce an opposite response. Indeed, upon application of ice water, ACLP^R60F09^ neurons demonstrated obvious increase in calcium signal indicated by GCAMP (Supplemental Figure 2). As temperature first dropped and then rose rapidly after the application of ice water, it was not clear if the calcium response was caused by temperature drop or rise. We then used the temperature controller to control temperature dropping monotonically from 27 °C to 18 °C without recovery. Decreasing the temperature from 27 °C to 18 °C within 30 seconds caused a noticeable response in larval ACLP^R60F09^ neurons, with an average peak response of about 30% increase in fluorescent intensity (Fig. 3e and 3f). ACLP^R60F09^ neurons’ response to temperature drop was completely abolished in presence of TTX, indicating that they receive cold signal from other neurons (Fig. 3e). Thus, ACLP^R60F09^ neurons could respond to both acute temperature rise and drop. In addition to these acute temperature changes, long term exposure to constant cool temperatures such as incubation at 18 °C for 18 hours could produce strong signals as visualized by Ca-LexA technique, while 18 hours of incubation at a warm temperature of 27 °C did not induce detectable signal (Supplemental Figure 3). These data together showed that ACLP^R60F09^ neurons could respond to both light and thermal stimulation. Taken together, integration of thermal signal and light signal in ACLP^R60F09^ neurons could be crucial for the negative regulation of larval light avoidance by temperature.

**Fig. 3.**
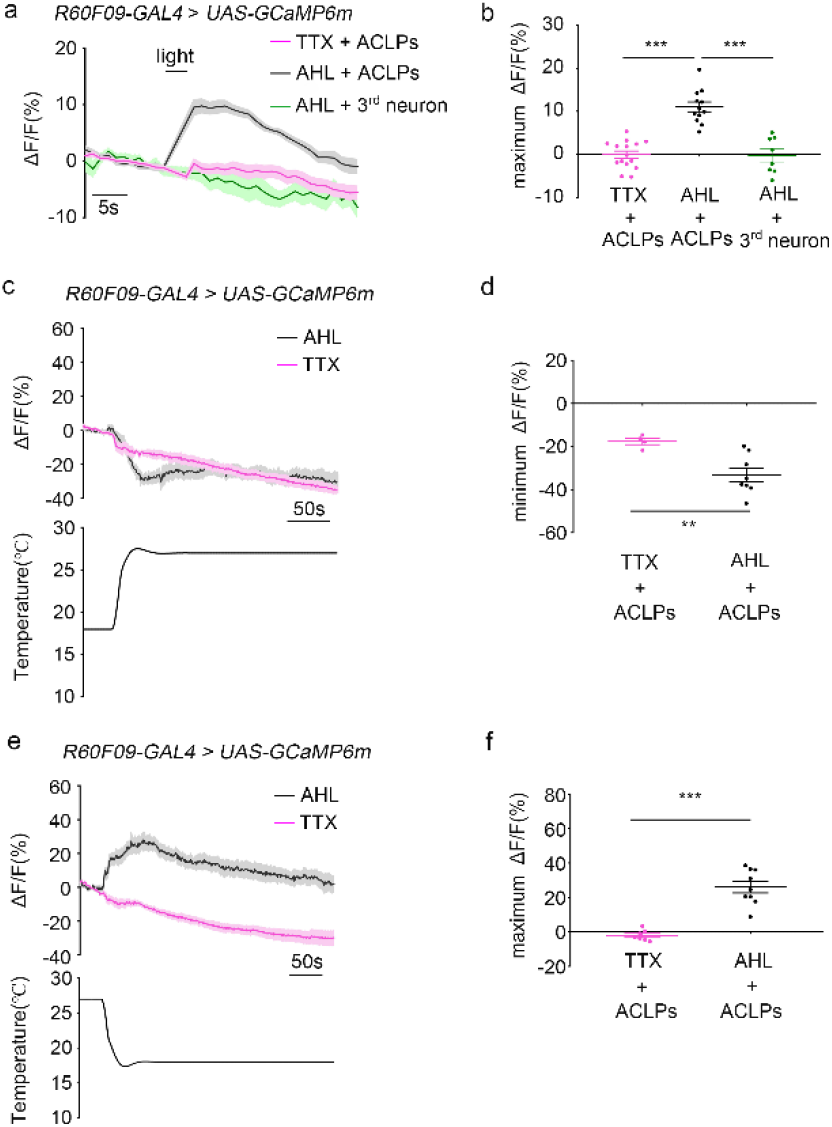
ACLP^R60F09^ neurons are responsive to both cold and light stimuli. (a) Ca^2+^ imaging of ACLP^R60F09^ neurons responses to 3 seconds of 460 nm blue-light stimulation. The 3^rd^ pair of brain hemisphere neurons labeled by *R60F09-Gal4* does not respond to light. Treating the neurons with 20 μM TTX for 4 minutes abolishes the response in ACLP^R60F09^ neurons. n=15 for TTX treated ACLP^R60F09^ neurons. n=12 for control ACLP^R60F09^ neurons treated with AHL. n=8 for the 3^rd^ pair of *R60F09-Gal4* labeled neurons treated with AHL. (b) Statistics of peak responses in (a). *** *P* < 0.001, unpaired *t*-test. Error bars, SEM. (c) Ca^2+^ imaging of ACLP^R60F09^ neurons’ responses to temperature rise from 18 °C to 27 °C. Treating the neurons with 20 μM TTX for 4 minutes abolishes the response. n = 4 for TTX treated ACLP^R60F09^ neurons. n = 8 for control ACLP^R60F09^ neurons treated with AHL. (d) Statistics of peak responses within 40 seconds after temperature change in (c). (e) Ca^2+^ imaging of ACLP^R60F09^ neurons’ responses to temperature drop from 27 °C to 18 °C. Treating the neurons with 20 μM TTX for 4 minutes abolishes the response. n = 7 for TTX treated ACLP^R60F09^ neurons. n = 9 for control ACLP^R60F09^ neurons treated with AHL. (f) Statistics of peak responses within 40 seconds after temperature change in (e). ** *P* < 0.01, *** *P* < 0.001, unpaired *t*-test. Error bars, SEMs.

### 4. *ACLP^R60F09^ neurons could be downstream to pdf*-LaNs *in visual pathway*

As the ACLP^R60F09^ neurons are morphologically close to *pdf-*positive lateral neurons (*pdf*-LaNs) which are known secondary visual pathway components (Supplemental Figure 4a), we wondered if ACLP^R60F09^ neurons received light signal from *pdf*-LaN neurons. We tested possible interactions between ACLP^R60F09^ neurons and *pdf*-LaNs using the GFP reconstitution across synaptic partners (GRASP) technique. Robust GRASP signal was observed between putative dendrites of ACLP^R60F09^ neurons and axonal termini of *pdf*-LaNs (Fig. 4a-d), suggesting that the ACLP^R60F09^ neurons might directly receive inputs from *pdf*-LaNs. To verify the existence of a functional connection, we combined optogenetics and calcium imaging to test whether directly exciting *pdf*-LaNs could activates ACLP^R60F09^ neurons. We expressed red light sensitive Chrimson in *pdf*-LaNs and GCAMP in ACLP^R60F09^neurons in larvae fed on food supplied with *trans*-retinal. Upon red light stimulation of *pdf*-LaNs, ACLP^R60F09^ neurons showed obvious calcium response, with an average peak response of more than 20% increase in fluorescent intensity (Fig. 4e and f). As *pdf*-LaNs can receive the light signal from larval photoreceptors (Larderet et al., 2017), it is thus possible that ACLP^R60F09^ neurons could receive light signal from *pdf*-LaNs.

**Fig. 4.**
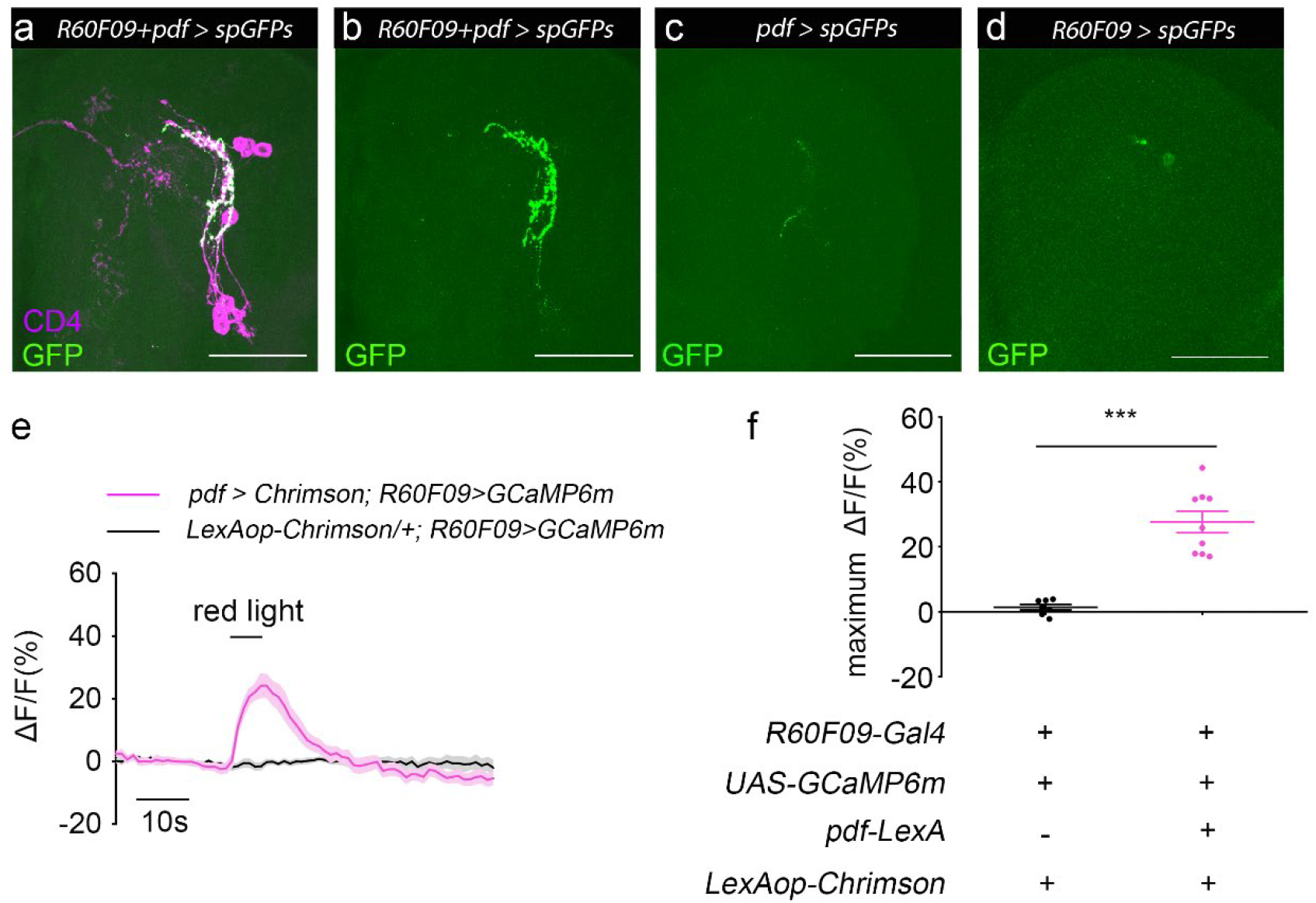
*pdf*-LaNs are upstream to ACLP^R60F09^ neurons. (a-d) ACLP^R60F09^ neurons and *pdf*-LaNs show strong GRASP signal (a and b) and no obvious detectable signal was seen in controls (c and d). GFP indicates GRASP signal in green; anti-CD4 indicates both ACLP^R60F09^ neurons and *pdf*-LaNs in magenta. Scale bars, 50 μm. (e) ACLP^R60F09^ neurons showed obvious Ca^2+^ response when *pdf*-LaNs expressing Chrimson were activated by 590 nm red light. n=9 for *pdf* > *Chrimson*; *R60F09* > *GCaMP6m*. n=7 for *Chrimson*/*+*; *R60F09* > *GCaMP6m*. (f) Quantification of peak Ca^2+^ response in (e). \*\*\**P* < 0.001. Error bars, SEMs.

### 5. *ACLP^R60F09^* neurons *could inhibit CLPN neurons*

As ACLP ^R60F09^ neurons appeared to be morphologically close to CLPN^R82B09^ neurons that were found to mediate the light induced head cast response in larval light avoidance (Supplemental Figure 4a and 4b), we wondered if ACLP^R60F09^ neurons could directly target on CLPN^R82B09^ neurons to affect light avoidance. We used GRASP technique to test possible interaction between ACLP^R60F09^ neurons and CLPN^R82B09^ neurons which could be labeled by promoter *GMR82B10*. ACLP^R60F09^ neurons and CLPN^R82B09^ neurons showed robust GRASP signal (Fig. 5a and 5b) while no signal was seen in the controls (Fig. 5c and 5d), suggesting that the ACLP^R60F09^ neurons might directly interact with CLPN^R82B09^ neurons. To further explore whether CLPN^R82B09^ neurons receive signal input from ACLP^R60F09^ neurons, we again combined optogenetics and calcium imaging to see if directly exciting ACLP^R60F09^ neurons could induce response in CLPN^R82B09^ neurons. When the ACLP^R60F09^ neurons expressing *LexAop-Chrimson* were excited by red-light, CLPN^R82B09^ neurons showed a maximal decrease of about 30% in calcium signal (Fig. 5e and 5f). This meant that CLPN^R82B09^ neurons received inhibitory signal input from ACLP^R60F09^ neurons. Indeed, when we co-stained *R60F09-Gal4* labeled neurons with the antibody against GABA, an inhibitory neurotransmitter, colocalization was seen in cell bodies of all the three *R60F09-Gal4* labeled neurons in brain hemispheres (Fig. 5g to 5r). This meant that ACLP^R60F09^ neurons were GABAergic. As CLPN^R82B09^ neurons had been identified to express a GABAA receptor RDL (Zhao et al., 2019), it was thus highly possible that ACLP^R60F09^ neurons inhibited CLPN^R82B09^ neurons through GABA/RDL interaction. Therefore, ACLP^R60F09^ neurons seemed to play inhibitory roles in larval light avoidance behavior.

**Fig. 5.**
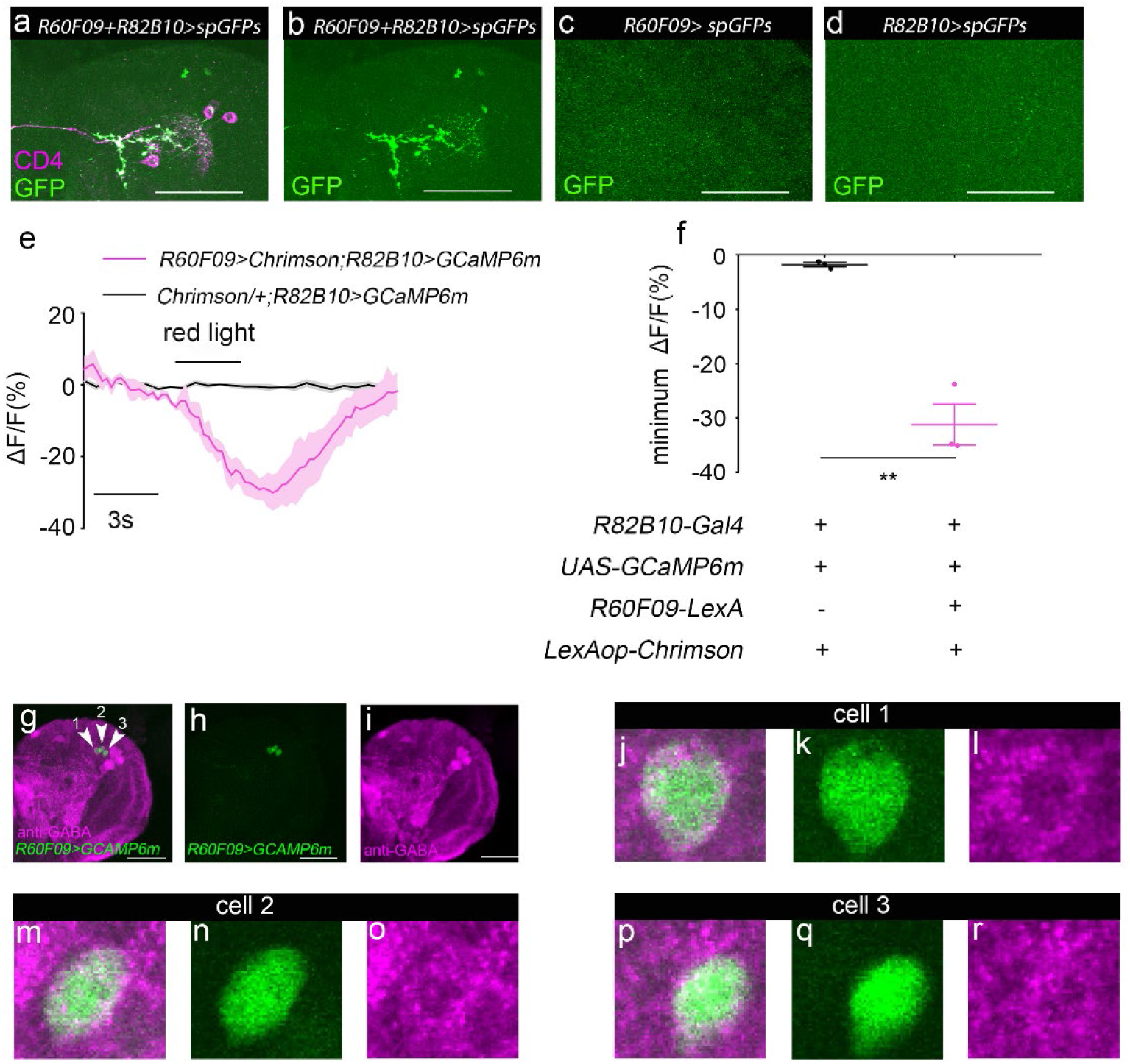
ACLP^R60F09^ neurons inhibit CLPN^R82B09^ neurons. (a-d) ACLP^R60F09^ neurons and CLPN^R82B09^ neurons show strong GRASP signal (a and b) and no obvious detectable signal was seen in controls (c and d). GFP indicates GRASP signal in green; anti-CD4 indicates both ACLP^R60F09^ neurons and CLPN^R82B09^ neurons in magenta. Scale bars, 50 μm. (e) Activation of ACLP^R60F09^ neurons inhibited CLPN^R82B09^ neurons. n = 3 for *R60F09 > Chrimson*; *R82B10 > GCaMP6m*. n = 3 for *Chrimson/+*; *R82B10 > GCaMP6m*. (f) Quantification of peak Ca^2+^ response in (e). ** *P* < 0.01). Error bars, SEMs. (g-r) ACLP^R60F09^ neurons are GABAergic. The three neurons in one brain hemisphere (g-i) are separately shown in (j-r). cell 1,2 and 3 are separately shown in (j-l), (m-o), and (p-r). anti-GABA signal is in magenta. GCAMP signal driven by *R60F09-Gal4* is in green. Arrowhead in (g) indicates the cell bodies of the three neurons. Note that *R82B10-Gal4* and *R82B10-LexA* were used to label CLPN^R82B09^ neurons. Scale bar is 50 μm in (g-i).

### 6. ACLP ^R60F09^s’ response to light was reduced at lower temperature

As ACLP^R60F09^ neurons were activated by temperature drop and inhibited by temperature rise, how could the inhibitory ACLP^R60F09^ neurons mediate the negative regulation of light avoidance by temperature, that is, enhanced light avoidance at lower temperature and repressed light avoidance at higher temperature? To explain this apparent paradox, we need to find out how light and thermal information were integrated in ACLP^R60F09^ neurons. We then tested the response of ACLP^R60F09^ neurons to light at 18 °C and 27 °C. Calcium responses of larvae expressing GCAMP in ACLP^R60F09^ neurons to 1-second light pulses at interval of 20 seconds at 18 °C and subsequently 27 °C were recorded. To reduce the effect of spontaneous activity on neuronal response, we used the average amplitudes of peak responses to the first five light pulses to measure the response amplitude. ACLP^R60F09^ neurons’ response to light stimulation decreased significantly at 18 °C compared to 27 °C (Fig. 6a and 6b), by a relative amplitude of more than 30% on average (Fig. 6c). As light response in ACLP^R60F09^ neurons was weaker at lower temperature, the light induced inhibition of ACLP^R60F09^ neurons on CLPN^R82B09^ neurons was thus relieved to allow stronger aversive reorientation response. These results could well explain the paradox and were in consistent with our previous observation that blocking ACLP^R60F09^ neurons abolished the thermal regulation of larval light avoidance.

**Fig. 6.**
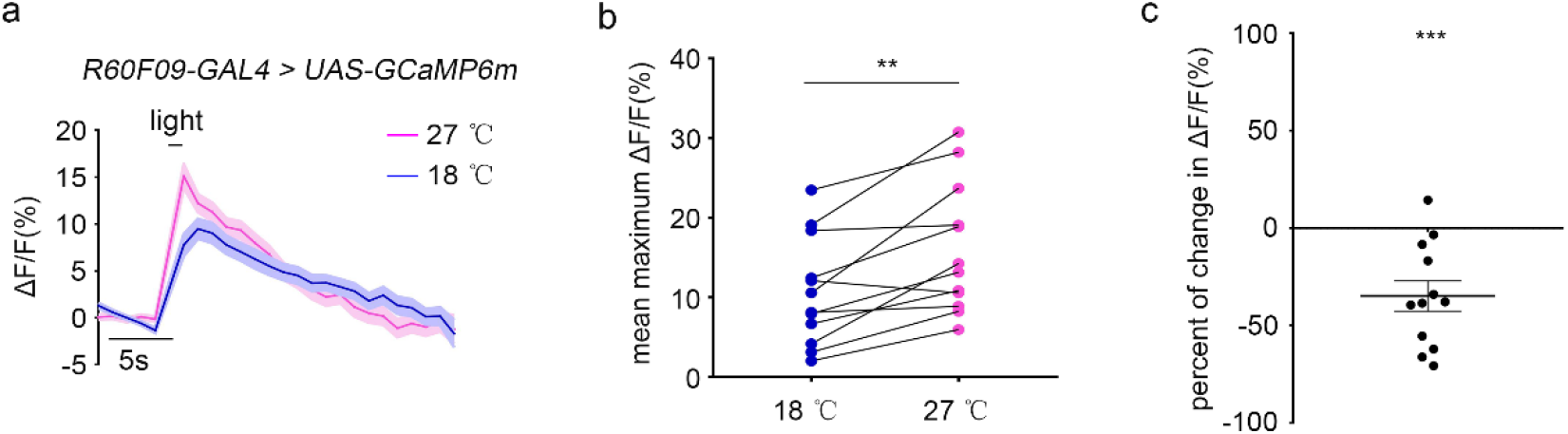
ACLP^R60F09^ neurons’ response to light was weaker at cool temperature than at warm temperature. (a) Ca^2+^ imaging of ACLP ^R60F09^ neurons’ responses to blue light at 18 °C and 27 °C. 460 nm light stimulation was applied for 1 second. n = 12. (b) Mean peak Ca^2+^ responses of ACLP^R60F09^ neurons to blue light at 18 °C and 27 °C. ** *P* < 0.01, paired *t*-test. n = 12. (c) The percent of decrease in the average amplitude of ACLP^R60F09^ neurons’ response to blue light at 18 °C compared with that at 27 °C. ** *P* < 0.01, *** *P* < 0.001, *t*-test against zero level. n = 12. Error bar, SEMs.

## Discussion

In this study, we found that *Drosophila* larval light avoidance was negatively regulated by temperature. When temperature dropped, the response of ACLP^R60F09^ neurons to light was decreased. As ACLP^R60F09^ neurons could inhibit the CLPN^R82B09^ neurons that command the light avoidance behavior, the reduction in ACLP^R60F09^ neurons’ light response at lower temperatures facilitated the light avoidance behavior by relieving the inhibition on CLPN^R82B09^ neurons.

The enhanced light avoidance at lower temperature should be good for survival of *Drosophila* larva. It is assumable that when temperature drops before arrival of winter, *Drosophila* larvae are more likely to avoid light and hide in shelter-like places. This will help adult fruit flies to overwinter in shelters such as ground holes or crevices (Izquierdo, 1991) which can protect them from being found by predators. The reduced general activity and metabolism at low temperatures further add to the importance of staying in shelter for overwintering.

In our results, despite that ACLP^R60F09^ neurons’ response to light was repressed at 18 °C, it was still quite significant. This meant that ACLP^R60F09^ neurons might still exert inhibition on CLPN^R82B09^ neurons. However, blocking ACLP^R60F09^ neurons did not change larval performance in light avoidance assay at 18 °C (Fig. 2). One possible explanation for such paradox could be that ACLP^R60F09^ neurons’ activity upon light stimulation at 18 °C was not high enough to substantially suppress the function of CLPN^R82B09^ neurons.

For explanation of ACLP^R60F09^ neurons’ function in the negative regulation of larval light avoidance by temperature, there are two remaining questions to be answered. The first question is from where do the ACLP^R60F09^ neurons received the cood and warmth signal, as ACLP^R60F09^ neurons do not sense temperature change by themselves. Cold sensing DOG and TOG neurons have been well characterized, but they all project directly into SOG area that is distant from ACLP^R60F09^ neurons. There must be some interneurons that transfer the cold signal onto ACLP^R60F09^ neurons. Sensory neurons for warmth in larva is so far not known (Barbagallo and Garrity, 2015). Another possibility is that ACLP^R60F09^ neurons receive thermal signals from other uncharacterized thermal sensing neurons. Neurons that express temperature-sensitive molecules such as various channels of the TRP family (Flockerzi 2007; Venkatachalam and Montell, 2007; Rosenzweig et al., 2008; Zhong et al.,2012; Fowler and Montell, 2013; Turner et al., 2016), molecules of the IR family (Li and Gong, 2017; Knecht et al.,2016; Budelli 2019), Brivido1-3 (Gallio et al., 2011) and rhodopsins (Shen et al., 2011) could be candidates. More work is required to fill the gap.

The second and probably more puzzling question is why ACLP^R60F09^ neurons’ response to light was reduced as temperature dropped. Since the light stimulated ACLP^R60F09^ neurons were excited by temperature drop and inhibited by temperature rise, it is straightforward to assume that light and low temperature facilitate each other’s excitatory effect on ACLP^R60F09^ neurons. Unexpectedly, the light response was repressed at lower temperature. This is probably because that lower temperature facilitates certain cellular event in ACLP^R60F09^ neurons that antagonizes the neurons’ response to light, or that higher temperature inhibits cellular event that facilitates the neurons’ response to light. Another explanation is that the integration of light and thermal signal occurs in some upstream neurons. In this case, the reduced light response at lower temperature in ACLP^R60F09^ neurons is a result of the upstream event.

In summary, our work disclosed a new behavioral paradigm of integration of temperature and light sensing and characterized the related neural basis. Future unraveling of the underlying mechanism will add to our understandings of cross-modal sensory integration in animal central brain.

## Acknowledgements

We thank J. Liu for constructive discussions, Y. Pan and F. Guo for sharing fly strains. We acknowledge the Bloomington *Drosophila* stock center and Qinghua *Drosophila* Stock center for providing the fly stocks and the core facilities of Medical School of Zhejiang University for technical support and reagents. This work is supported by the National Natural Science Foundation of China (31070944, 31271147, 31471063, 31671074 and 61572433), and the Fundamental Research Funds for the Central Universities, China (2017FZA7003).

## Methods

### Fly strains

Most flies were reared at 25 °C on standard culture medium under 12h:12h light/dark cycles. Flies used for optogenetic experimental were raised on food supplemented with 0.2 mM all-trans-retinal in constant darkness at 25 °C. The following strains were used in this work: *w^1118^*, *WT-CS*, *R60F09-GAL4* (BL39255), *R82B10-GAL4* (BL46717), *R60F09-lexA* (BL61576), *pdf-LexA*, *UAS-TNTG* (BL28838), *UAS-GCaMP6m* (BL42748,42750), *UAS-mCD8-GFP* (BL5137), *UAS-CD4::GFP1-10*, *hsFLP, UAS-FRT-stop-FRT-Chrimson, UAS-mLexA-VP16-NFAT* (Li and Gong, 2015), *LexAop-CD4::GFP11*, *LexAop-Chrimson* (BL55138,55139). Except for *R60F09-GAL4* and *R60F09-lexA,* all other strains can be referred to Zhao et al., 2019 unless specifically mentioned.

### Larval light avoidance assay

The procedure of light spot assay was largely same as previously described (Zhao et al., 2019). In short, half of the petri dish containing a 1.5% agar plate was covered to create a dark environment, white light above the petri dish illuminate the uncovered half. Light intensity of 550 lux corresponded to 23.3 μW/mm^2^, 170 lux corresponded to 9.1μW/mm^2^ at maximal readings (S401C, Thorlabs, Inc). Twenty 3^rd^ instar larvae were placed on the agar plate to choose between light and dark for different time (5min or 10min or 15min or 20min) at different temperatures (15°C, 18°C, 20°C, 23°C, 25°C and 27°C). Light preference index (PI) was calculated as: PI = (number of larvae in the dark half–number of larvae in the light half)/(number of larvae in the dark half + number of larvae in the light half).

### Calcium imaging

Calcium imaging was done using 2^nd^ or 3^rd^ instar larvae as in previous report (Zhao et al., 2019). If 3^rd^ instar larvae were used, they were dissected in AHL (Adult Hemolymph-Like) solution to remove the posterior part and keep the central brain exposed, and then transferred into a chamber formed by reinforcing rings on a cover glass and covered with another cover glass. If 2^nd^ instar larvae were used, the single larva was directly covered with a cover glass without reinforcing ring. The cell bodies of neurons were imaged. Ca^2+^ imaging was performed with Olympus FV-1000 two-photon microscope with 40X water objective lens at room temperature of 23 °C. Infrared laser at 910 nm was used for illuminating the GCaMP6m. For calcium imaging response of ACLP ^R60F09^ neurons to controlled cooling (from 27 °C to 18 °C) and warming (from 18 °C to 27 °C) or to light, the 2^nd^ instar larva or dissected 3^rd^ instar larva between cover glasses was placed on the aluminum plate surface of a custom-made temperature controller. Precise cooling and warming were controlled by the temperature controller. The light stimulus was 460nm blue light at intensity of about 28.076 μW/mm^2^. For calcium imaging response of ACLP^R60F09^ neurons to ice water, 200μl ice water was added on the glass slide.

For calcium imaging response of pharmacologically isolated ACLP^R60F09^ neurons to light and to thermal stimuli, larvae were dissected and soaked in AHL (adult hemolymph like) saline containing 20μM TTX for 4 minutes before recording the ACLP^R60F09^ neurons’ calcium response. The soaking solution was AHL for control samples.

For Ca^2+^ imaging ACLP ^R60F09^ neurons’ responses to optogenetic activation of *pdf*-LaNs and CLPN^R82B09^ neurons’ responses to optogenetic activation of ACLP^R60F09^ neurons, eggs of proper genotypes were laid on food supplied with 0.2mM trans-retinal and raised at 25°C in constant darkness until 3^rd^ instar larvae. 590nm red light was used to activate *pdf*-LaNs or ACLP^R60F09^ neurons. Images were acquired at 1.109s per frame at resolution of 512 × 512 in pixels.

For quantitative analysis of Ca^2+^ imaging data, we used ImageJ (imageJ.nih.gov/ij) to batch process images to determine fluorescence intensity of regions of interest. We subjectively selected a few sequential images right before thermal stimulation to calculate average fluorescence intensity (F) as the basal level. Changes in fluorescence intensity (ΔF) were calculated and ΔF/F was used to denote Ca^2+^ responses.

### Ca-LexA (calcium-dependent nuclear import of LexA) imaging

Eggs of proper genotypes were laid on food and raised at 25°C under 12h:12 h light/dark cycles for two days. The larvae were then divided into two groups and placed in incubators with internal temperature of 18 °C and 27 °C respectively. After 18 hours, larvae were dissected, fixed, and imaged with confocal microscopy (see following).

### Immunochemistry and Confocal Microscopy

We dissected brains from 3^rd^ instar larvae in PBS, fixed in PFA (PBS containing 4% paraformaldehyde) for 60min at room temperature, washed three times for 0.5h each time in PBT (PBS containing 0.5% Triton X-100), and blocked for 1 h in PBT containing 5% goat serum. The samples were then incubated with primary antibodies (mouse anti-PDF, 1:100, PDF-C7 concentrate, DSHB; rabbit anti-CD4, 1:200, Cat. ab133616, Abcam; rabbit anti-GABA, 1:50, Cat. A2052, Sigma-Aldrich) overnight at 4 °C, followed by three times (0.5h each time) wash in PBT. The samples were then incubated with secondary antibody (Alexa 647 goat anti-mouse, 1:100, Cat. A21235, Thermo Fisher; Alexa 647 goat anti-rabbit, 1:100, Cat. A27040, Thermo Fisher; Alexa Fluor 594 goat anti-rabbit, 1:100, Cat. 33112ES60, YESEN) for 2 h at room temperature and washed in PBT three times for 10min each time in darkness. Finally, brains were mounted and viewed. Images were acquired using an Olympus FV1000 confocal laser scanning microscope with 20X-, 40X oil- or 60X oil-objective lens at resolution of 1024 × 1024 in pixels.

### Single neuronal morphology labeling experiment

Eggs of proper genotypes were laid and collected on food. The food containing the collected eggs was evenly dispersed in a plastic petri dish, then the petri dish was placed in a 37 °C water bath for 5 minutes, after which the food containing the eggs was transferred to a vial containing standard culture medium. Eggs were reared at 25 °C under 12h:12 h light/dark cycles for 3 days. 3^rd^ instar larvae were dissected, fixed, and imaged.

### Statistics

For all the tests, paired *t*-test, unpaired *t*-test or one-way ANOVA with Tukey’s post hoc test were used. Error bars in scatter plot and shaded areas flanking curves represented SEM.

---

**Supplementary Figure 1.**
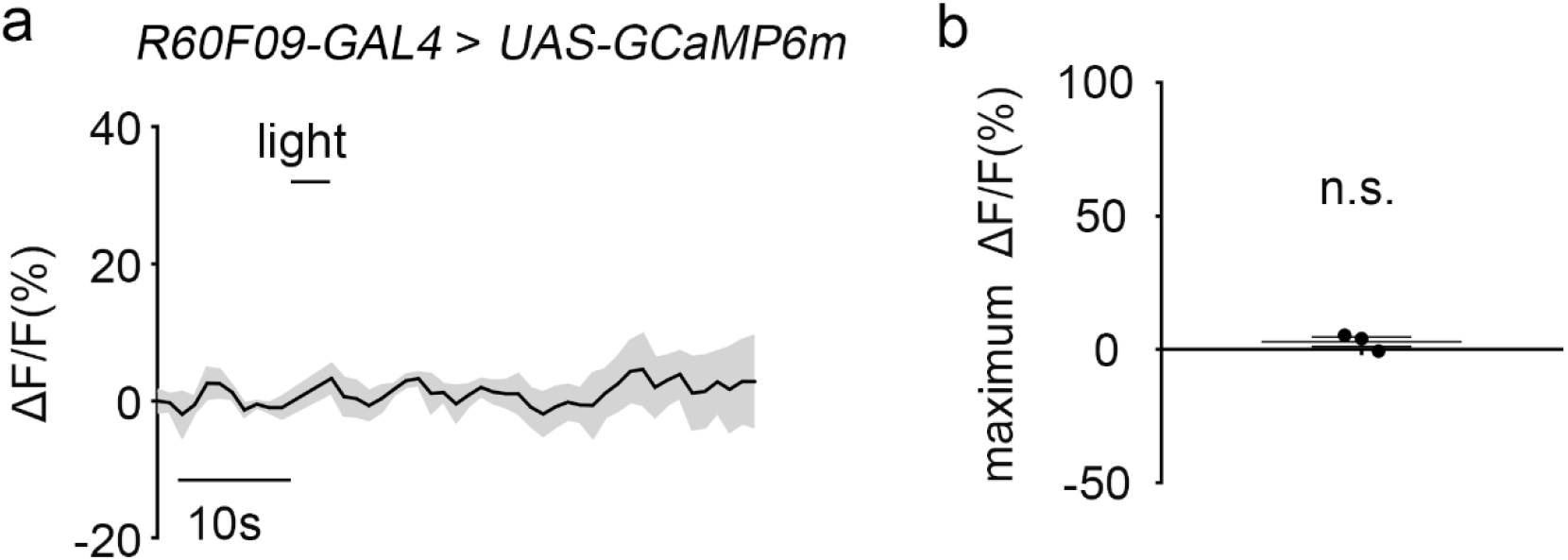
Neurons in the suboesophageal ganglion (SOG) labeled by *R60F09-GAL4* do not respond to light stimulation. (a) Ca^2+^ responses of neurons in the suboesophageal ganglion (SOG) labeled by *R60F09-GAL4* to blue light. 460 nm blue-light stimulation was applied for 3 seconds. n = 3. (b) Statistics of peak response in (a). n.s., *P* > 0.05, *t*-test against zero level. Error bars, SEMs.

**Supplementary Figure 2.**
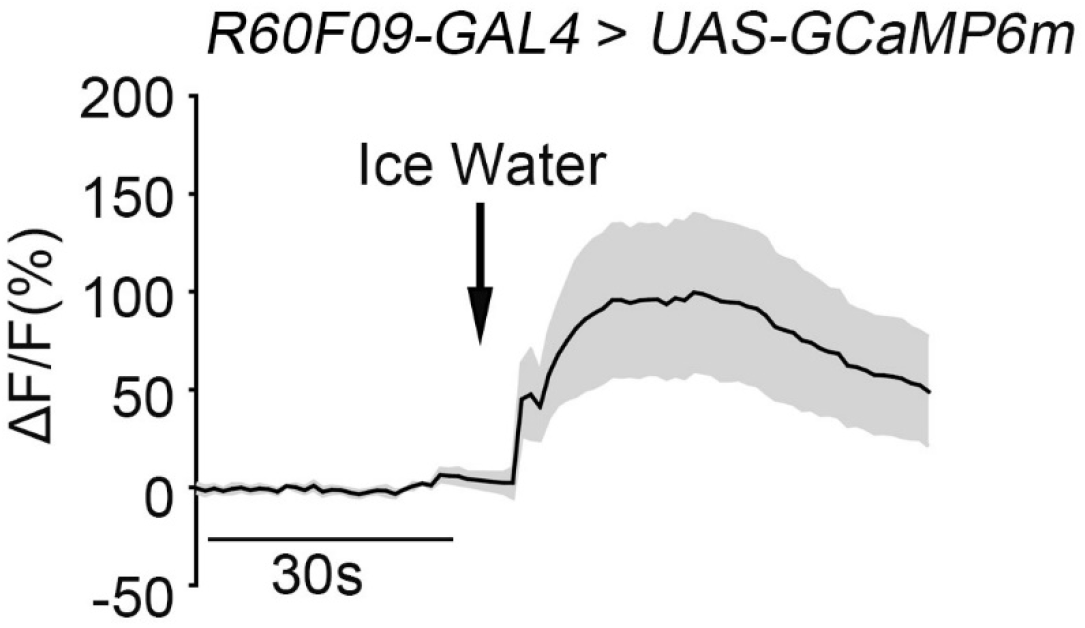
Ca^2+^ imaging of ACLP^R60F09^ neurons’ responses to addition of 200 μl ice water. n = 7.

**Supplementary Figure 3.**
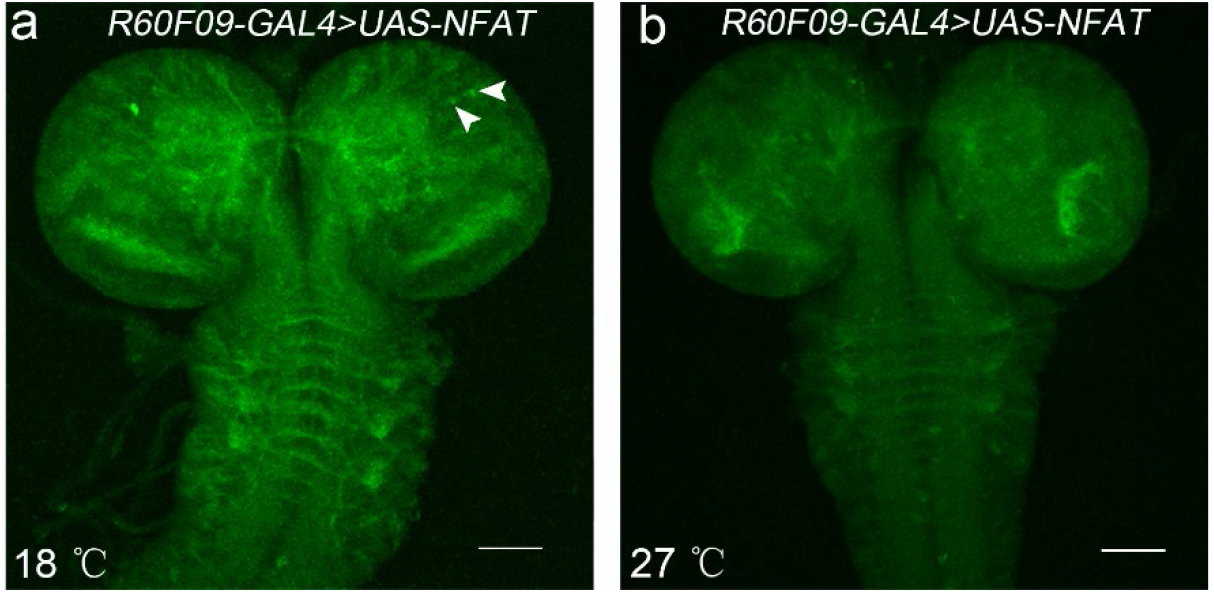
ACLP^R60F09^ neurons respond to long-term cool and warm temperature. (a, b) ACLP^R60F09^ neurons are labeled after 18-hour cold stimulation at 18 °C (a), while were not labeled after 18-hour warm stimulation at 27 °C (b). Arrowheads indicate the signals in neuronal cell bodies. Scale bars, 50 μm.

**Supplementary Figure 4.**
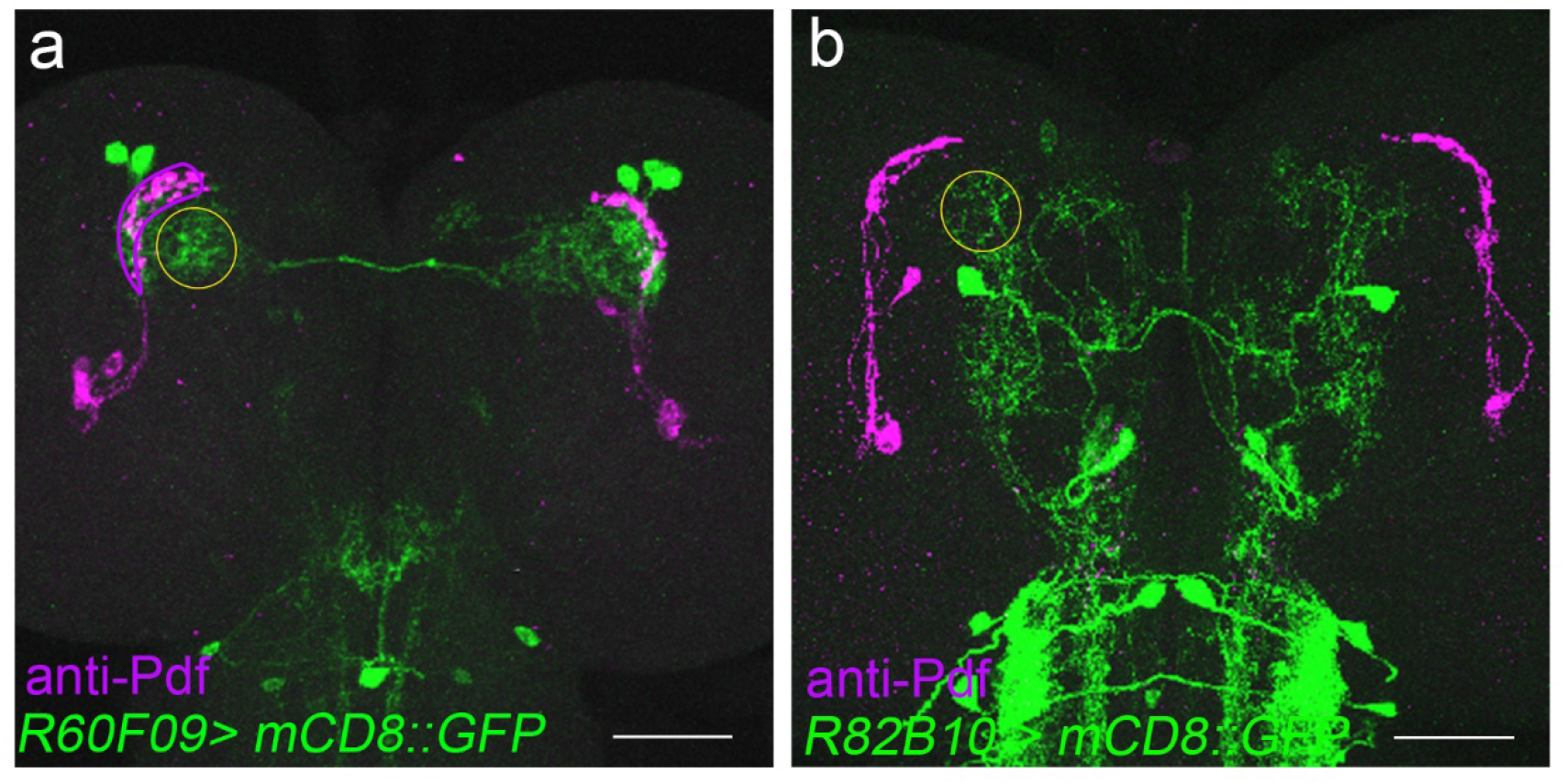
Co-staining of ACLP^R60F09^ neurons (a) and CLPN^R82B09^ neurons (b) with anti-Pdf in larval brain hemispheres. According to the locations relative to that of the *pdf*-LaNs, neurites of ACLP^R60F09^ neurons and CLPN^R82B09^ neurons are close to each other. GFP signal indicates the neurons in green, anti-Pdf signal indicates the morphology of *pdf*-LaNs in magenta. The yellow circles outline the putative overlapping region between dendrites of ACLP^R60F09^ neurons and CLPN^R82B09^ neurons. Scale bars, 50 μm.

